# Metabolic plasticity increases evolvability and drives persistence to PARP inhibitor in ovarian cancer

**DOI:** 10.1101/2025.10.08.681266

**Authors:** Adriana Del Pino Herrera, Joon-Hyun Song, Karla Torres-Arciga, Susoma Halder, Kadin El Bakkouri, Jowana Obeid, William M. Burke, Mehdi Damaghi, Meghan C. Ferrall-Fairbanks

## Abstract

PARP inhibitors (PARPi) are the standard maintenance therapy for ovarian cancer, but the frequent emergence of resistance remains a major clinical challenge. Emerging evidence suggests that tumor cell plasticity is an important mechanism enabling this adaptive resistance. In this study, we combined both experimental and computational techniques to model this evolutionary process by long term treatment of the OVCAR3 ovarian cancer cell line with Olaparib (a PARPi). We then performed single-cell RNA sequencing (scRNAseq) on both naïve and persistent cells to capture cancer cells heterogenous response. scRNAseq profiling suggested an adaptive metabolic reprogramming in response to long term Olaparib treatment. We then scored single cells for metabolic function, and persister cells were found to have reprogrammed their glutamine metabolism. To test these findings, both persister and naïve cells were cultured in glutamine-deprived media and persister cells were found to have higher viability than the naïve cells particularly when exposed to Olaparib. To test the cells capacity to switch their metabolic state toward glycolysis, we performed Seahorse mitochondrial and glycolytic stress tests. These analyses revealed that persister cells were less dependent on glutamine metabolism and instead exhibited increased glycolytic capacity compared to naïve cells. This metabolic shift resulted in an acquired dependency on glucose, representing a potential therapeutic vulnerability. Together, these results indicate that prolonged Olaparib exposure promotes metabolic plasticity in ovarian cancer cells. To explore how this plasticity might be exploited therapeutically, we developed a proof-of-concept mathematical model that leverages metabolic state transitions to optimize treatment scheduling.

**ONE SENTENCE SUMMARY:** Ovarian cancer cells exploit metabolic plasticity to increase their evolvability to persist PARP inhibitor treatment.

## INTRODUCTION

Ovarian cancer is the fifth deadliest cancer in the United States, accounting for 20,890 patients diagnosed and 12,730 deaths from the disease in 2025 [1]. Difficulty in ovarian cancer diagnosis is due to its unremarkable symptoms, where over 70% of cases are diagnosed at advanced stages of disease when it has already metastasized [2]. This has led to the death of 1 in 6 patients within the first three months of diagnosis and a 5-year survival rate of 51.6% [1]. Current treatment strategies involve a combination of surgical resection and chemotherapy with platinum-based chemotherapeutic and taxane-based agents, which are assumed to be effective at the onsets of the disease for all patients. If the cancer metastasized, neoadjuvant chemotherapies are administered before surgical resection [3]. These chemotherapeutic agents are delivered systemically, which leads to severe toxicities in the patients. In hopes to decrease toxicity, poly ADP-ribose polymerase inhibitors (PARPis) are used as maintenance therapy [4]. The PARP enzyme fixes single-strand DNA (ssDNA) damages. PARPis bind to PARP, making it unable to repair ssDNA damages leading to double-strand breakages in the DNA, ultimately resulting in cell death. However, PARP is not the only enzyme able to repair DNA damages in cells [5]. A process called homologous recombination led by BRCA1/2 can repair double-strand damages to ensure cell viability [6]. PARPis, such as Olaparib, are widely used as maintenance therapy for ovarian cancer patients, particularly those harboring BRCA1/2 mutations, but they are also used in some BRCA-wild-type tumors with homologous recombination deficiency (HRD) [7]. The effectiveness of PARPi therapy has been shown in clinical trials, where the risk of recurrence on BRCA-deficient ovarian cancer patients was reduced by 40-70%, and half of the patients experienced a 5-year progression-free survival [8].

Independent of BRCA status, not all patients respond positively to PARPis and most eventually acquire resistance to the treatment [4]. More than 40% of patients with BRCA1/2 mutations do not respond to PARPis, and several patients will develop resistance after prolonged administration to these therapies [9]. Similar to other cancers, ovarian cancer cells eventually evolve resistance to targeted therapy and thus combination therapies should be considered [4], [10], [11]. Increasing evidence suggests that tumor cell plasticity plays a critical role in evolving resistance to cancer, allowing cancer cells to dynamically reprogram transcriptional, metabolic, and DNA-repair pathways in response to therapeutic stress [12]–[14]. Here, we show that metabolic plasticity enables ovarian cancer cells to persist under PARP inhibition on their evolutionary trajectory towards full resistance. Temporary resistance, or persistence, was developed in the OVCAR3 ovarian cancer cell line with a commonly used PARPi, Olaparib. While OVCAR3 cells do not carry BRCA1/2 mutations, they lack the homologous recombination machinery to repair breakages in DNA making them an ideal cell system to study PARPi effects [15], [16]. Single-cell transcriptomics profiling of naïve and persister OVCAR3 cells, showed a metabolically reprogrammed persister state that increased cell fitness by the upregulation of glycolytic capacity and glutamine metabolism, suggesting a shift in nutrient utilization under therapeutic stress. Metabolic flux analyses further reveal a compensatory increase in glycolytic capacity, uncovering a shift toward glucose-dependent energy production that represents a potential therapeutic vulnerability. Functional perturbation confirms that persister cells maintain higher viability under glutamine deprivation, indicating reduced dependence on glutamine metabolism. Together, our results identify metabolic plasticity as a key adaptive strategy enabling evolving evolvability to resistance to Olaparib. For the first time we discovered one mechanism of increased evolvability through metabolic switching and increased glucose metabolism reserve. We suggest that targeting dynamic metabolic dependencies could provide a framework for evolution-informed combination therapies to delay or prevent resistance in ovarian cancer. Integrating these experimental findings with population dynamics modeling, we demonstrate that exploiting metabolic state transitions through adaptive treatment scheduling may limit the expansion of persistent populations.

## RESULTS

### Olaparib persistence phenotype described at single-cell resolution in OVCAR3

To investigate the mechanisms of PARPis persistence in ovarian cancer, we performed scRNAseq on two populations of the BRCA1/2 wildtype ovarian cancer cell line OVCAR3: untreated naïve cells and cells that had evolved persistence following prolonged (more than two months) exposure to Olaparib at 10 μM (**Fig. 1A**). We used the term persistence over resistance because even after 6 months of Olaparib exposure, when Olaparib was removed from the media the cells reverted back to their parental drug sensitive phenotype [2]. After persistent lines were developed, cell viabilities for the parental (naïve) and persister cell populations were assessed using Cell Counting Kit-8 (CCK-8) viability assay (n=3) at different treatment concentrations (0, 5, 10, 25, 50, 100 μM Olaparib). Naïve cells showed higher sensitivity to PARPi at drug concentration of 10uM. At 50, and 100 μM Olaparib treatment conditions, persister cells had higher viability implying the development of the drug tolerance (**Fig. 1B**). This observation is in line with recent studies showing increased adaptability of cancer cells to PARPi [17].

**Figure 1.**
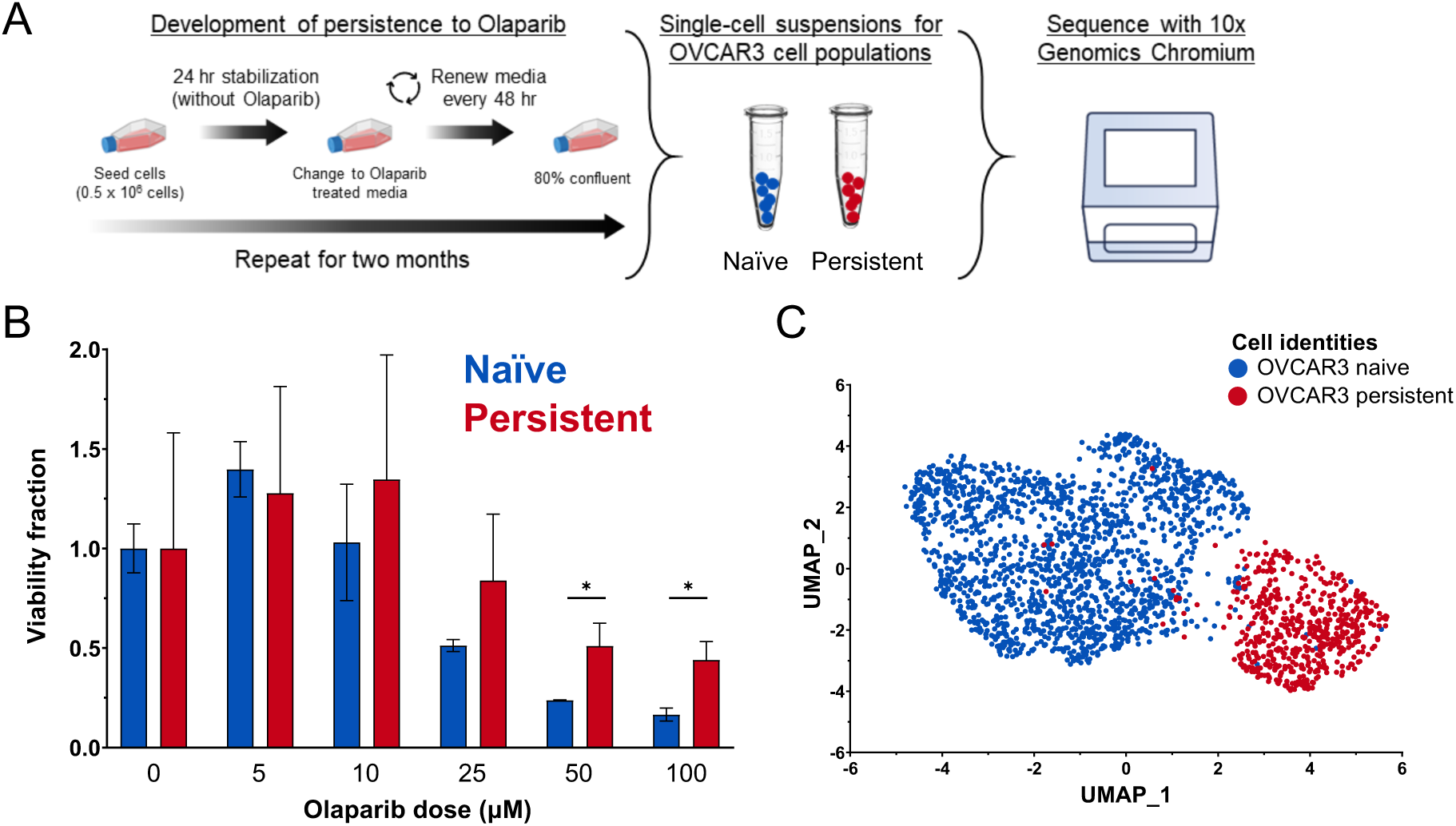
Divergent evolutionary trajectory captured by Olaparib persistence in OVCAR3 at single-cell resolution. **A**, Schematic of the cell culturing process and development of Olaparib persistence. OVCAR3 cells were cultured in 10μM Olaparib treated media for two months until persistence evolved. Single-cell suspensions were transcriptomically sequenced using the 10x Genomic Chromium platform. **B**, A one-week CCK-8 clonogenic assay shows cell viability fractions as low as 0.1659 for OVCAR3 naïve populations (n=3) at higher Olaparib doses. The persistent population shows a 2-fold larger cell viability fraction of 0.4397 at high Olaparib doses compared to the parental cell lines (n=3). **C**, UMAP visualization of single cell RNA-sequencing data labeled by cell type (naïve cells, n=1712, blue and persistent cells, n=550, red).

Passage number matching naïve and persister cells were then sequenced at a single-cell resolution using the 10X Genomics Chromium technology to identify differentially expressed genes and metabolic pathways characteristic of each population [18], [19]. A total of 2,348 cells were sequenced (1,766 naïve and 582 persistent) and filtered to remove low-quality cells. The cell number after filtering was reduced to 2,262 (96%), which were then log-normalized and scaled before dimension reduction was performed using principal component analysis. Graph-based clustering was used to identify phenotypically defined clusters and visualized using UMAP. Based on cell identity, the naïve and persistent populations have two distinct areas implying their distinct evolved biological signature (**Fig. 1C**).

### Metabolic signature analysis of OVCAR3 naïve and persistent populations indicate glutamine dependency

We performed clustering using the Louvain algorithm and characterized five distinct phenotypic clusters within naïve and persistent populations (**Fig. 2A**). To understand the single-cell transcriptomic diversity in the dataset, the differentially expressed genes for each of the five distinct clusters were identified (**Supplementary Tables S1-5**). The significant and upregulated genes (p-value > 0.05 and average log2 fold-change) were used to identify the characteristic Reactome metabolic pathways and gene markers for each cluster (**Fig. 2A and C and Supplementary Tables S6-10**). Additionally, all the clusters contained mixed populations of both naïve and persister cells, suggesting that the development of persistence might be a continuous process. Clusters 0, 3 and 4 expressed a higher proportion of naïve cells (98%, 99%, and 98%, respectively) whereas cluster 1 was 98% composed of persister cells (**Fig. 2B**). Cluster 0 was mostly characterized by pathways focused on the translation of proteins with the upregulation of genes such as AIMP1, MRPL16 and SRP72. Given that translation is a pathway commonly performed by the majority of the cells, the expression of these markers across the clusters was similar. Cluster 1 (98% persister cells) showed increased expression of respiratory electron transport markers (MT-ND3 and TIMM21) indicating a potential energy switch towards oxidative phosphorylation in the persistent population. Clusters 2, 3, and 4 which are composed of a majority of naïve cells showed an upregulation of the cell cycle mitotic, Rho GTPase cycle and DNA repair pathways, respectively. Some characteristic markers of these clusters were cyclins and kinases (CCNB1 and PLK1) in C2, genes encoding for structural proteins (SPTAN1 and ACTG1) in C3 and check point inhibitors as well as DNA repair genes (CLSPN, FEN1, and RMI2) in C4. The upregulation cell cycle related genes and pathways in naïve cells suggests an ideal target for PARPi treatment. Specific Gene Ontology terms were also identified per cluster, which highlighted specific functional mechanisms incorporated in the pathways of translation, respiratory cell transport, cell cycle mitotic, Rho GTPase cycle, and DNA repair for C0, C1, C2, C3, and C4, respectively (**Supplementary Tables S11-15**).

**Figure 2.**
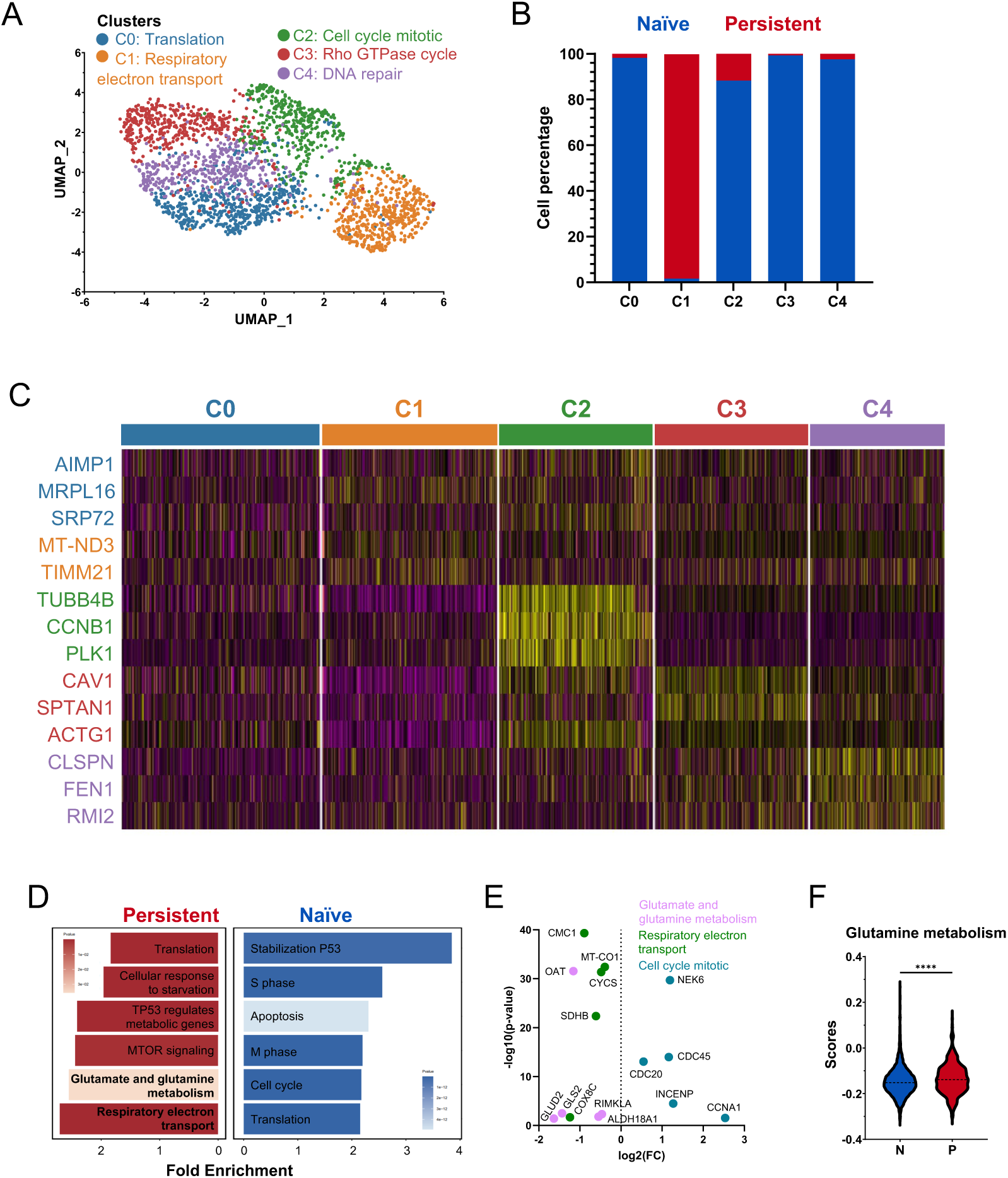
Transcriptomic differences between naïve and persistent populations indicate persistence dependence towards the glutamine metabolism. **A,** Graph-based clustering identified 5 distinct clusters from the single-cell data, each characterized by a Reactome pathway. Clusters 0, 2, 3, and 4 were composed of mostly naïve cells and characterized by translation, cell cycle mitotic, Rho GTPase cycle and DNA repair pathways, respectively. **B**, Bar-graph with naïve and persistent cell proportions per clusters. Cluster 1 contains 98% persistent cells while clusters 0, 2, 3, and 4 contain between 88-99% naïve cells. **C**, Heatmap of significant marker genes (p-value < 0.05) for each cluster corresponding to the characteristic pathways. **D**, Top naïve and persistent Reactome pathways identified using significant upregulated differentially expressed genes for each group (p-value <0.05, average log2(fold-change) > 0). The most enriched pathways for naïve and persistent populations were the stabilization of P53 and the respiratory electron transport, respectively. **E**, Scatter plot showing the differential expressed genes in naïve cells compared to persistent for three pathways of interest. **F**, After evaluating 42 unique metabolic pathways and calculating a pathway score for each cell in the dataset, the glutamine metabolism expressed significant difference between naïve and persistent populations (p<0.0001), using the Mann-Whitney test.

Focusing on transcriptomic differences between naive and persister cells, the significant and upregulated (p-value > 0.05 and average log2 fold-change) markers for each population were also identified (**Supplementary Tables S16-17**). These gene lists were used to identify characteristic naive and persistent Reactome pathways (**Fig. 2D**). Metabolic pathways with the highest fold enrichment (FE) and significant p-values were shown on the bar plot (lists for all the Reactome pathways can be found in **Supplementary Tables S18-19**). Generally, naïve cells revealed an upregulation of cell cycle-related pathways such as S phase (FE: 2.56, p-value < 0.0001), M phase (FE: 2.2, p-value <0.0001), and cell cycle mitotic (FE: 2.17, p-value < 0.0001). When focusing on differentially expressed genes for naïve compared persister cells (**Fig. 2E**), kinases, cyclins and genes encoding for cell cycle regulatory proteins (NEK6, CCNA1, CDC45, and CDC20) showed a significant upregulation in naïve cells. The upregulation of cell cycle pathways and apoptosis (FE: 2.3, p-value < 0.0001) confirmed the effectiveness of PARPi therapies at inducing cell cycle arrest and ultimately cell death in naïve cells [20]. Additionally, naïve OVCAR3 cells upregulate the stabilization of P53 (FE: 3.85, p-value < 0.0001) while TP53 regulates metabolic genes in persister cells (FE: 2.41, p-value < 0.0001) [21]. OVCAR3 is characterized by the loss of the tumor-suppressor gene P53 leading to survival of damaged cells, aggressive tumor growth and resistance [15]. In the context of OVCAR3 naive cells, the stabilization of this protein would allow for a less aggressive phenotype and treatment effectiveness compared to persister cells where TP53 controls metabolic processes indicating unresponsiveness to treatment. A study testing the reactivators P53 showed that these have a synergistic effect with Olaparib enhancing the efficacy of PARPi in OVCAR3 cells [22]. The persistent population also showed an upregulation of the respiratory electron transport pathway (FE: 2.7, p-value < 0.0001) which, similar to cluster 1, these cells prioritize energy production through oxidative phosphorylation. While most cancer cells perform glycolysis to obtain energy also known as the Warburg effect, OVCAR3 Olaparib-persister cells under no treatment showed a switch towards aerobic oxidative phosphorylation [23]. Recent studies have shown that silencing the oxidative phosphorylation pathway can re-sensitize PARPi resistant cells in ovarian cancer patient derived models [24].

Another metabolic pathway to highlight was the glutamine and glutamate metabolism which was upregulated in persister cells (FE: 2.56, p-value < 0.05). The respiratory electron transport and glutamine metabolism results were coupled with the downregulation of mitochondrial genes (CMC1 and COXC8) and amino acid metabolism genes (OAT, GLUD2, and GLS2) in naïve cells compared to persister. This indicates that persister cells upregulate their glutamine production as part of their persistent mechanism to Olaparib. Persister cells also revealed an upregulation of metabolic pathways such as mTOR signalling (FE: 2.44, p-value < 0.05) and the regulation of metabolic genes by switch in persister cell growth. Lastly, the expression of 1,456 genes linked to 42 KEGG [25] metabolic pathways was evaluated to test the hypothesis that persistence may be related to glutamine dependence. A score was calculated for each cell based on the expression of the gene list associated with each pathway. The glutamine metabolism scores showed a significant difference (p-value < 0.0001) between the naïve and persistent populations (**Fig. 2F**). Additional RNA velocity analysis using scVelo and DeepVelo was used to evaluate transcriptomic changes between states (clusters) [26]–[28]. The velocity vectors indicated that cells in clusters 0, 2, 3, and 4 have the potential to transition to a transcriptomic state similar to cells in cluster 1, from a naïve state to a persistent state (**Supplementary Methods and Supplementary Figure S1**). This transition was evaluated in terms of glutamine metabolism gene expression across the latent time and showed that GLS (responsible for the first reaction in the glutamine metabolism) appeared towards the end of the latent time. Specific Gene Ontology terms per naive and persistent populations were also identified which highlighted specific functional mechanisms (**Supplementary Tables S20-21**). Overall, these results indicate that reprogramming glutamine metabolism plays a major role in evolving persistence to Olaparib in OVCAR3 cells [6], [29].

### OVCAR3 persistent cells increased glycolytic capacity and metabolic plasticity to Olaparib treatment

To further investigate glutamine metabolism regulation in response to Olaparib treatment, both naïve and persister cells were cultured in media with 0 μM Olaparib (untreated) with and without glutamine (Gln+ and Gln− respectively) and 50 μM Olaparib (treated) with and without glutamine (Gln+ and Gln− respectively). CCK-8 assays were performed to determine the effects of glutamine deprivation with and without Olaparib treatment on cell viability after one week of treatment (n=3). The viability of the naive cells decreased by 35% when they were treated with Olaparib and Gln+, while the viability of the persistent population remained 100% under those conditions (**Fig. 3 A and B**). When glutamine was depleted from the media in absence of Olaparib, the viability dropped more dramatically in naïve cells compared to persister (by 72% versus 48%) against 0 μM Olaparib/Gln+ media, implying the glutamine dependency of naïve cells and losing this dependency during the term of adaptation to Olaparib. However, under Olaparib treatment in glutamine-deprived media, the viability of naïve cells slightly dropped by 8%, while that of the persistent populations stayed the same, suggesting that under Olaparib treatment both naïve and persister cells are less dependent on glutamine metabolism. This fast metabolic switch in both cells confirms the cancer cells metabolic plasticity. Also, higher viability in the persistent population suggests that cells could be upregulating their intrinsic glutamine production to withstand the effects of the drug and the lack of glutamine in the environment. These results led to the hypothesis that, upon PARPi treatment, cancer cells prioritize glutamine for DNA repair over energy production. To test whether cancer cells shift their metabolism toward glycolysis, a Seahorse analysis was performed to measure cells glycolytic phenotype compared to their mitochondrial respiration under Olaparib treatment with and without glutamine. Under no treatment and glutamine-added conditions both naive and persister cells showed high oxygen consumption rates (OCR) suggesting that both populations obtain energy under aerobic conditions. Both naïve and persister cells treated with Olaparib in the presence of glutamine have increased extracellular acidification rate (ECAR) implying the increased glycolysis and switching to glucose metabolism. Interestingly, glutamine deprivation decreases both naïve and persister cells mitochondrial respiration and glycolysis dramatically. However, when treated with Olaparib the glycolysis is increased to basal level. This metabolic plasticity persisted in persister cells, which maintained higher glycolysis levels even in the absence of treatment (**Fig. 3C and 3D**). Analysis of the oxygen consumption rate (OCR) indicates a decrease upon treatment with PARP inhibitors in both naïve and persister cells, corroborating the ECAR data and suggesting a metabolic shift towards increased glycolysis (**Fig. 3E**).

**Figure 3.**
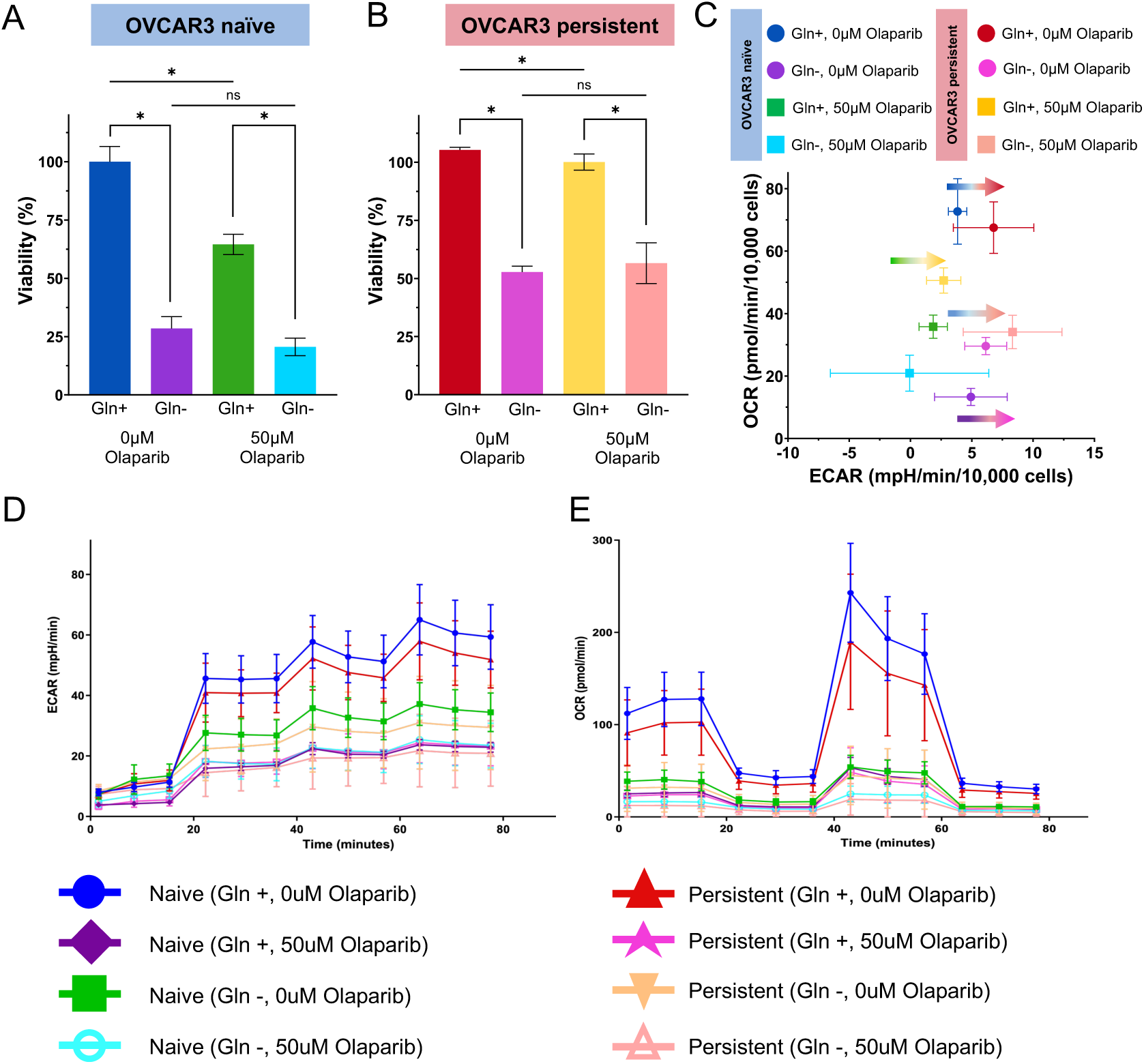
OVCAR3 persistent cells demonstrate high viability under Olaparib treated and glutamine deprived conditions. **A,** Viability assay of OVCA3 naïve cells treated with Olaparib in absence or presence of Gln. **B**, Viability assay of OVCA3 persistent cells treated with Olaparib in absence or presence of Gln. Viability is measured using CCK8 kit and data is the results of 3 biological replicates with 4 technical replicates in each biological replicates. Student t-test is used for analysis. **C**, OCR/ECAR analysis of the naïve and persistent OVCAR3 cells from Seahorse assay. Arrows show the transition toward more glycolysis under PARPi treatment. **D**, Extracellular acidification rate (ECAR) of OVCA3 cells in conditions similar to **A**. **E**, Oxygen consumption rate (OCR) of OVCA3 cells.

### Mathematical modelling simulations of naïve and persistent cells show a decrease in cumulative drug dosages for glutamine deprivation and adaptive therapy schedules compared to continuous therapy schedules

Mechanistic mathematical modelling was used to study cell growth dynamics over time under different environmental conditions (0μM Olaparib, 20μM Olaparib, Gln+, and Gln− conditions). Cell counts were collected every 48-72 hours for 772 hours (n=5-18) (**Supplemental Figure S2**) and were cultured in well plates, which constrain exponential growth in a density-dependent manner. Therefore, a logistic growth model was selected to model the growth dynamics of these populations *in vitro*. Logistic growth assumes a dependence of population growth rate on population density [2]. When the total population reaches a carrying capacity then it will stop growing. Logistic growth is described as:

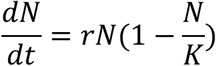

where *N(t)* is the population size at time *t*, *r* is the cell growth rate (in hours^−1^), and *K* is the carrying capacity. Cell counts were normalized by the counts on day 0 to have a better representation of population dynamics across all conditions. These counts were used to estimate logistic growth rates (r) and carrying capacities (K) in Julia by minimizing the sum of squared errors between experimental data and the model. Estimated logistic growth parameters for each cell population (OVCAR3 naïve and persistent) were computed under 0μM Olaparib/Gln+ environmental conditions (**Supplemental Table T22**). In terms of growth rates, persistent cells present a higher growth rate than naïve cells. However, naïve cells have a higher carrying capacity than persistent cells indicating that under standard media conditions (0μM Olaparib/Gln+) naïve cells will dominate in the population.

The growth rates and carrying capacities estimated for naïve and persistent cells under untreated and glutamine added conditions were used to create co-culture simulations between these populations across the different media conditions and different initial persistence percentage. This co-culture model was described using the following equations:

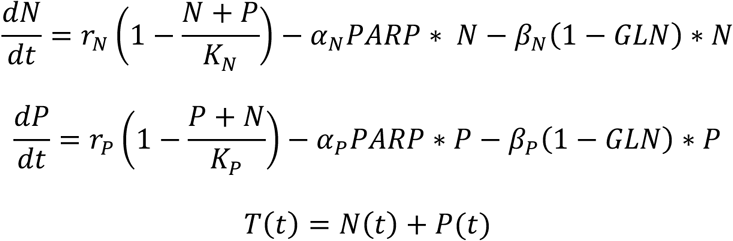

where *N(t)* and *P(t)* are the naïve and persistent population sizes at time *t*, *r_N_* and *r_p_* are the cell growth rate (in hours^−1^), and *K_N_* and *K_p_* are the carrying capacities for the naïve and persistent population previously estimated under untreated and glutamine added conditions. The effect of treatment and glutamine deprivation was modelled as an on and off term that would change based on the magnitude of the factors of *α_N_* and *α_p_* for

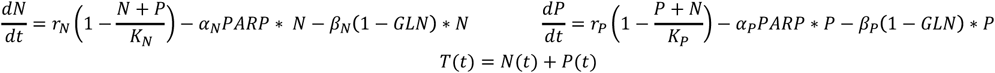

treatment effect and *β*_*N*_ and *β*_*P*_ for glutamine deprivation. These terms were estimated for naïve and persistent populations individually by fitting the above equations, which contained the growth rates and carrying capacities under untreated and glutamine added conditions, to the data collected using different media conditions. For instance, different *α*_*N*_ and *α*_*P*_ where calculated for glutamine added and glutamine deprived media conditions. Similarly, different *β*_*N*_ and *β*_*P*_ were calculated for both untreated and treated conditions. Generally, *α*_*N*_ > *α*_*P*_ both under glutamine added and glutamine deprived media which is consistent with previous results of Olaparib greatly (**Supplementary Table S23**) affecting the viability of naïve cells while maintaining viability of the persistent group. Additionally, *β*_*N*_ > *β*_*P*_ both under untreated and treated conditions indicating that glutamine deprivation affects the naïve population more. This is reflected in the co-culture simulations (**Figure 4**). Co-culture simulations under four different continuous media conditions (0μM Olaparib/Gln+, 0μM Olaparib/Gln−, 20μM Olaparib/Gln+, and 20μM Olaparib/Gln−) and three different initial persistence percentages (25%, 50%, and 75%) were initially created across 768 hours. Under these four continuous media conditions and 25% initial persistence, naïve cells initially dominate in the population. However, naïve cells are only dominant by 768 under standard media conditions (0μM Olaparib/Gln+) and 25% initial persistence. Focusing on the total population growth, glutamine deprivation under untreated conditions (0μM Olaparib/Gln−) causes a 0.64-fold decrease in the total population by 768 hours across all initial persistence percentages compared to standard conditions (0μM Olaparib/Gln+). Olaparib treatment alone (20μM Olaparib/Gln+) only causes a 0.13-fold decrease in the total population compared to standard conditions. However, when both glutamine deprivation and treatment are combined (20μM Olaparib/Gln−) then the total population experiences a decrease of 0.85-fold compared to standard conditions. This indicated that the combination of glutamine deprivation and Olaparib treatment at the tumor site can achieve a greater cancer population size reduction than treatment alone. In order to achieve this decrease in the total population, a cumulative drug dose of 340μM Olaparib (20μM delivered every 48 hours) is necessary. This cumulative dosage will result in several patient toxicities and complications in treatment.

**Figure 4.**
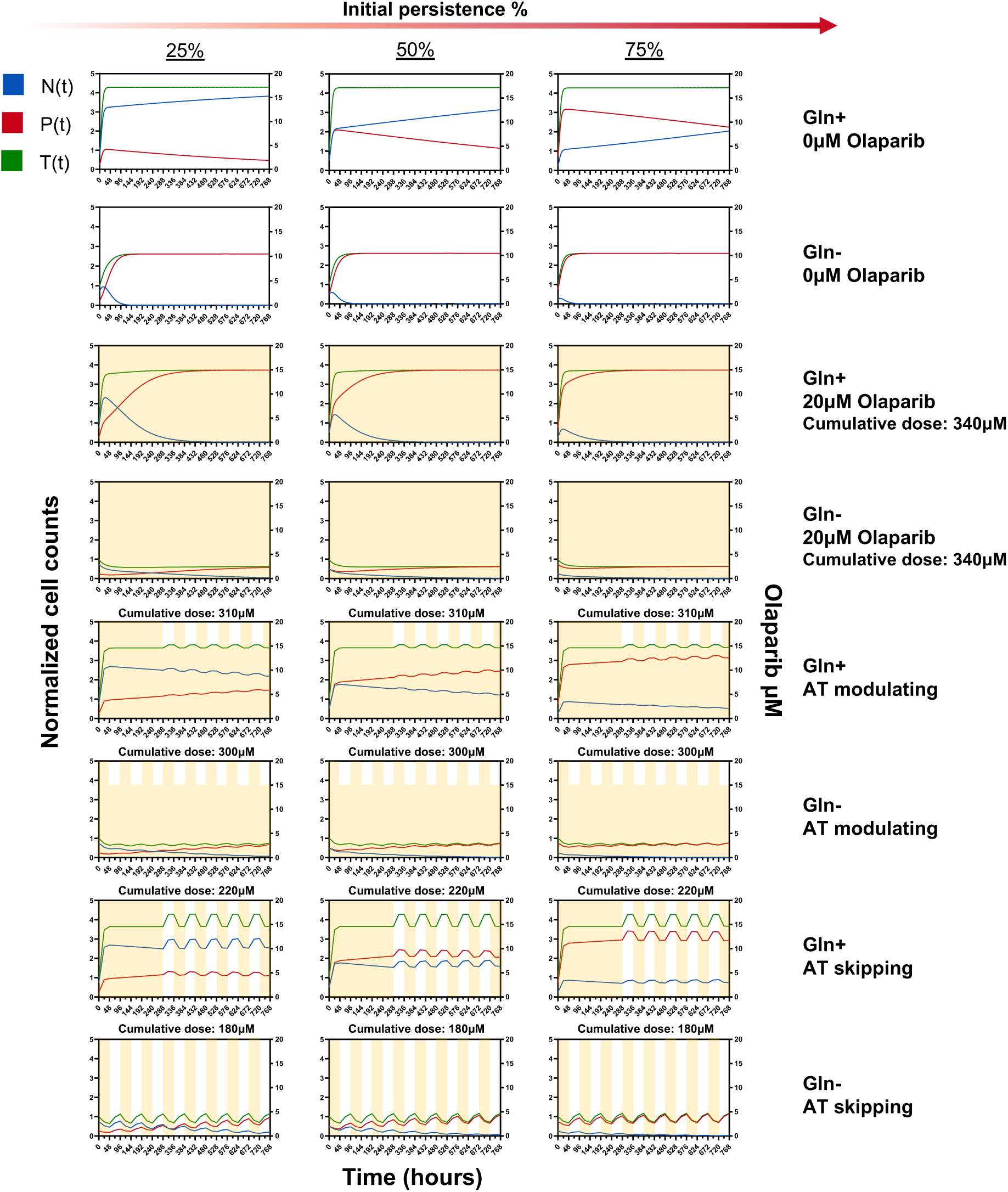
Mathematical models showed different growth dynamics of OVCAR3 naïve and persistent cells under different environmental conditions. OVCAR3 naïve and OVCAR3 persistent growth dynamics were estimated logistic growth curves computed on the normalized cell counts. Growth rates and carrying capacities for both OVCAR3 naïve and OVCAR3 persistent cells were parameterized in Julia using growth data under untreated and glutamine-added conditions (0μM Olaparib/Gln+). Using data from all other media conditions (0μM Olaparib/Gln−, 50μM Olaparib/Gln+, 50μM Olaparib/Gln−), an effect of treatment and glutamine deprivation linear terms were estimated. The resulting parameters were used to create co-culture simulations show the effects of different media conditions on the growth dynamics of sensitive (blue), resistant (red) and total (green) populations. The different initial persistence percentage shows that sensitive cells will dominate in the population only the initial persistence is 25% and the media is untreated, and glutamine is added. While treatment alone and glutamine deprivation alone each contribute to a decrease in the total population, this reduction is further amplified when glutamine deprivation and Olaparib are applied. Adaptive therapy simulation show that evolutionary treatment delivery reduces the cumulative drug dosage compared to continuous drug delivery. Different adaptive therapy schedules (AT modulating and AT skipping) were tested under glutamine added and deprived conditions. AT skipping under glutamine deprived conditions achieved the same treatment efficiency as the continues delivery of treatment with a much lower cumulative treatment dose.

To minimize cumulative drug dosages, in the addition to the four continuous media conditions already simulated, two different adaptive treatment schedules were also tested similar to Strobl et al. schedules [2]. For both schedules, the glutamine was either present or deprived (Gln+ or Gln− respectively). The first schedule (AT modulating) consisted in dynamically adjusting the drug dosage based on changed in the total population size. If the population changed by 10% in 48 hours then the drug dosage would be adjusted by 5μM. For example, if the total population size decreased by 10% then the drug dosage would decrease by 5μM (until it reached 0μM). If the population later increased by 10% then the drug dosage would increase by 5μM, reaching a maximum drug concentration of 20μM. The second schedule (AT skipping) consisted on skipping treatment periods if there was a total population reduction of 10% in a 48-hour period. Both schedules began by treating the population at an Olaparib concentration of 20μM and either modulating or skipping a dose (setting the treatment to 0μM) when the population decreased by 10%. For both schedules, glutamine deprived conditions (Gln−) showed a reduction in cumulative drug dosages. The lowest cumulative drug dosage was achieved through the glutamine deprived adaptive therapy skipping schedule (180μM cumulative dose). Compared to the continuous glutamine deprived and treated conditions (20μM Olaparib/Gln−), AT Skipping/Gln− results in a 0.63-fold increase in the total population by 768 hours while nearly utilizing nearly half the amount of Olaparib. Therefore, adaptive therapy is able to achieve similar results to continuous treatment in terms of total population size but using much lower values of drug. Glutamine deprivation and adaptive treatment has the potential to reduce treatment toxicities while achieving tumor control.

## DISCUSSION

Ovarian cancer is the second most lethal gynecological disease characterized by high tumor heterogeneity. The nature of the disease increases treatment challenges resulting in a high probability of recurrence even to targeted therapies such as PARPi. The mechanisms underlying resistance to PARP inhibition are not fully understood and are likely multifactorial. Known resistance mechanisms include restoration of homologous recombination, stabilization of replication forks, drug efflux, and alterations in PARP trapping efficiency [4]. However, these genetic mechanisms do not fully explain the rapid emergence of drug-tolerant populations observed in many tumors. Increasing evidence suggests that non-genetic adaptive processes, including cellular plasticity, play a critical role in enabling tumor cells to survive therapeutic stress [12]–[14]. Plasticity allows cancer cells to transiently reprogram transcriptional, epigenetic, and metabolic states, creating phenotypic diversity within the tumor population that can facilitate survival under drug pressure. In this study, we provide evidence that metabolic plasticity is a key adaptive mechanism enabling ovarian cancer cells to persist under Olaparib treatment (**Figure 1**). Using single-cell transcriptomic profiling of OVCAR3 cells exposed to prolonged Olaparib treatment, we identified the emergence of a metabolically distinct persister population. These cells exhibit transcriptional signatures consistent with altered nutrient utilization and metabolic rewiring. In particular, our analyses revealed changes in pathways related to glutamine metabolism and glycolysis, suggesting a shift in metabolic strategy under sustained PARPi pressure (**Fig. 2D-F and Figure 3**).

Glutamine plays a crucial role in the body by maintaining and promoting cell functions such as protein synthesis and serving as a precursor for nucleotide and neurotransmitter synthesis [30], [31]. In the context of cancer, the glutamine metabolism is often dysregulated and a GL2 overexpression indicates poor survival in cancers such as colon, blood, and ovarian [32]. Previous research has shown that glutamine depletion has avoided the development of resistance to chemotherapeutics in cancers such as pancreatic and liver cancer [31], [33]. In these cancer cell populations, cells upregulated their glutathione synthesis, the successor to the glutamine metabolism, to protect themselves from the effects of chemotherapy. The hypothesis that PARPi treatment causes cancer cells to shift glutamine metabolism toward DNA repair rather than energy production highlights an intriguing adaptive mechanism in cancer cell survival. PARP inhibitors, like Olaparib, target DNA damage repair pathways by inhibiting the PARP enzyme, which is crucial for repairing single-strand breaks in DNA [34]. This puts significant stress on the cancer cells’ ability to manage DNA damage. In response, cancer cells may alter their metabolic pathways, such as reallocating resources like glutamine, a versatile amino acid that plays a role in both energy production and biosynthetic pathways [35]. The hypothesis suggests that after PARPis treatment, glutamine’s primary role may shift toward aiding DNA repair rather than fueling the tricarboxylic acid (TCA) cycle for energy generation. This reallocation likely reflects the cells’ need to prioritize survival under DNA damage stress. These cancer cells do this adaptation by synchronizing with adjusting their cell cycle as also implicated in our single-cell data. These cells under Olaparib reduce their proliferation rate significantly and use the glutamine mainly for DNA repair rather than energy. One-week cell viability experiments under different media conditions combining the presence and absence of Olaparib and glutamine, showed that the viability of persister cells is not as affected under treated and glutamine deprived media compared to the untreated and glutamine deprived media (**Fig. 3A and B**). Whereas the viability of naïve cells decreased when treatment was added to glutamine deprived media. To further explore this, Seahorse analysis was used to measure metabolic shifts, particularly glycolysis (**Fig. 3C-D**). Functional experiments confirmed that these transcriptional changes correspond to altered metabolic dependencies. Persister cells displayed reduced reliance on glutamine metabolism and maintained higher viability than naïve cells under glutamine deprivation, particularly during Olaparib treatment. At the same time, metabolic flux measurements revealed a compensatory increase in glycolytic capacity, indicating that persister cells shift toward glucose-dependent energy production. This metabolic reprogramming likely provides a flexible energy supply that supports survival under therapeutic stress. Importantly, this shift also creates a potential metabolic vulnerability, as persister cells become increasingly dependent on glycolytic metabolism.

From an evolutionary perspective, this metabolic transition can be interpreted as a phenotypic adaptation that increases cellular fitness in the presence of PARP inhibition. Rather than immediately acquiring genetic resistance, cancer cells first adopt a metabolically permissive state that allows them to tolerate drug exposure. Such transient states are increasingly recognized as drug-tolerant persister states, which serve as intermediates along the evolutionary path toward stable resistance. If therapeutic pressure is sustained, these adaptive states may eventually become genetically or epigenetically stabilized, effectively “hard-wiring” the resistant phenotype into the population. Our findings therefore support a model in which metabolic plasticity increases the evolvability of cancer cells under therapy. By dynamically shifting between metabolic programs, cancer cells can explore alternative fitness landscapes and survive otherwise lethal conditions. This concept is consistent with broader evolutionary theories of cancer, which propose that phenotypic plasticity can accelerate adaptation by increasing the range of accessible phenotypes within a population.

An important implication of our findings is that metabolic plasticity may function as a mechanism that increases the evolvability of cancer cells under therapeutic pressure. Evolvability refers to the capacity of biological systems to generate heritable phenotypic variation that can be selected upon. In the context of cancer, non-genetic mechanisms such as transcriptional and metabolic plasticity can rapidly expand the phenotypic repertoire of tumor cells without requiring immediate genetic mutations. This expanded phenotypic landscape enables cancer populations to explore adaptive states that increase survival under treatment. Our data suggest that PARPi exposure selects for cells capable of dynamically reprogramming their metabolism, allowing them to occupy a metabolic state that is less vulnerable to drug-induced stress. Over time, these transient adaptive states may become stabilized through genetic or epigenetic alterations, effectively converting short-term persistence into stable resistance.

Importantly, the metabolic rewiring observed here suggests opportunities for therapeutic intervention. Because persister cells display **i**ncreased glycolytic capacity and altered nutrient dependencies, metabolic pathways may represent tractable targets for combination therapies with PARPis. Targeting glycolysis or glucose uptake, for example, may selectively impair the survival of persister populations while sparing naïve cells that rely more heavily on alternative metabolic pathways.

The inhibition of both glycolysis and glutamine metabolism in a breast cancer model showed that this phenomena could trigger an immune response and inhibit the growth and metastasis of tumors [36]. Thus, targeting glycolysis and glutamine metabolism in conjunction with PARP inhibition might increase the therapeutic efficacy by preventing persister cell survival, which is often a major hurdle in durable cancer treatment outcomes. However, we suggest that these inhibitions should consider the evolvability of the cancer cells and should be applied in an evolutionary designed manner [4]. For that reason, we built a mathematical model to simulate naïve and persistent cell growth under different combinations of Olaparib and glutamine (**Figure 4**). While a lower growth rate was estimated for naïve cells compared to persister cells, these cells presented a higher carrying capacity indicating naïve cell dominance in the population when combined with persister cells. When evaluating the effects of treatment and glutamine deprivation in these populations, parameter estimation showed that the growth of naïve cells is more negatively affected under treated or glutamine deprived conditions compared to persister cells. This suggests that persistent populations are intrinsically upregulating their glutamine metabolism to be able to enhance resistance. When evaluating co-culture simulations of naïve and persistent populations, it was observed that both the effects of treatment and the effects of glutamine deprivation affect the growth of the naïve population more negatively compared to persistent. However, these simulations show the potential effectiveness of the combination of Olaparib and glutamine deprivation. While total population decrease is observed in both continuous and adaptive therapy schedules, adaptive therapy is able to achieve a much lower cumulative drug dosage. In this scenario, adaptive therapy has the potential of achieving the control of the total population as well as the ability to greatly decrease the cumulative drug dosage across the treatment schedule. Moreover, adaptive therapy combined with glutamine deprivation has the potential to not only provide tumor control but to decrease treatment toxicities in patients since cumulative drug dosages would be much lower than continuous treatment schedules. The combination of data science, experimental, and mathematical modeling techniques have allowed to study the development of PARPi persistence in OVCAR3 through different angles which have pointed towards PARPi persistence plasticity towards glycolysis and glutamine metabolism. Further studies applied to multiple ovarian cancer cell lines and the expansion to animal models would validate this metabolic plasticity under PARPi treatments.

## METHODS

### Cell culture

OVCAR3 cells were acquired from American Type Culture Collection (ATCC) and cultured in RPMI medium (ThermoFisher) supplemented with 10% Fetal Bovine Serum and 1% penicillin/streptomycin. At all times cells were kept at 37°C and in 5% CO_2_. Cells were tested for mycoplasma contamination and were authenticated before the study started.

### Viability analysis (CCK-8 assay)

To measure cancer cells growth in the absence and presence of Olaparib treatment, cells were seeded in a 48 well flat-bottom plate and let them attach overnight. The media was replaced with treated growth medium, containing 0, 1, 10, 25, 50 or 100 μM Olaparib (AstraZeneca), and cell growth was monitored for 8 days. Viability assays were measured using CCK8 kit for at least 4 technical and 3 biological replicates.

### Cell growth dynamics

Cell growth was monitored by imaging the cells at 4X and 10X resolution using Cytation X (Agilent) time-lapse microscopy system. Confluency was measured based on bright field images and analyzed using the Gen5 Software.

### Single-cell RNA-sequencing

Chromium Single Cell 3′ Library, Gel Bead & Multiplex Kit and Chip Kit (10x Genomics) was used to encapsulate and barcode for cDNA preparation of OVCAR3 parental (naïve) and persistent cells. The libraries were constructed according to the manufacturer’s protocol, sequenced on an Illumina NovaSeq and mapped to the human genome (GRCh38) using CellRanger (10x Genomics).

Raw gene expression matrices generated per sample using CellRanger (v.3.0.1) were combined in R (v.4.2.0) and converted to a Seurat object using the Seurat R package [37]. Naïve and persistent Seurat objects were merged and studied in combination for further downstream analysis. Initially, low quality cells were filtered out of the dataset. Cells with unique feature counts under 200 and above the mean count reads plus two standard deviations were filtered from the dataset. Cells that expressed a high mitochondrial gene expression above the average expression for each cell and under 200 were also filtered. After filtering, the data was log-normalized, and the top 4287 number of highly variable features were identified using the FindVariableFeatures() function. The data was scaled to regress out variation due to differences in count and percentage of mitochondrial content. Then, a linear dimension reduction was performed using the highly variable features (PCA). A graph-based clustering approach (Louvain algorithm) was used to identify phenotypic clusters, which resulted in 5 distinct clusters composed of both naïve and persistent cells. Data was visualized using UMAP projects [37].

### Differential gene expression and pathways analysis

Differentially expressed genes for the dataset were identified using the Seurat FindMarkers() function. To identify differentially expressed genes on each cluster, the cluster number was input as the identity whereas to identify differentially expressed genes on the naive or persistent populations, these were input as the identity. Once the differentially expressed genes were identified for the population of interest, the significant and upregulated genes were filtered using p-value < 0.05 and average log2(fold-change) > 0 and converted to ENTREZ IDs. Both gene ontology (GO) and Reactome pathways databases from the Molecular Signatures Database (MSigDB) [38] were downloaded to Rstudio. These were then used as a reference to determine pathways characteristic of each population based on their significant and upregulated gene list compared to a background list of all the genes in the genes in the dataset. While GO terms provided functional mechanisms, Reactome pathways provided a focused metabolic view that was preferable for this study. Both p-values and fold enrichment values were considered when choosing characteristic pathways for each population.

### Metabolic pathway scoring

A list of 1,456 genes linked 37 KEGG and 5 Reactome metabolic pathways was used for gene enrichment analysis. The gene lists for each pathway were scored across each cell using the AddModuleScore() function. Scores for naïve and persistent populations were compared for each pathway using the Mann-Whitney U test.

### Seahorse XF Analysis

Glycolytic rate of OVCA3 and selected OVCA3 cancer cells was measured using Seahorse XF96 extracellular flux analyzer and a MST kit (Seahorse Biosciences). OCR and ECAR of cancer cells were determined by seeding them on XF96 microplates in RPMI medium until cells reached over 90% confluence. In these studies, seeding started with 20,000 cells (80% of well area). Measurements were determined 24 h later when the cells reached the 90% confluence. One hour before the Seahorse measurements, culture media were removed, and cells were washed with PBS followed by addition of base medium (nonbuffered Dulbecco’s Modified Eagle Medium supplemented with 25 mM glucose) or our nonbuffered only-glucose-containing solution. For glycolytic rate measurements, mitochondria inhibitors including rotenone (1 μM) and antimycin A (1 μM) were injected after basal measurements of ECAR and OCR of the cells under treatment to stop the mitochondrial acidification. Finally, data were normalized for cell number in each well from imaging by CytationX. Seahorse measurements were performed with four to six technical replicates, and these experiments were repeated four times.

Solutions for seahorse experiments. A total of 2 mM Hepes, 2 mM MES, 5.3 mM KCl, 5.6 mM NaPhosphate, 11 mM glucose, 133 mM NaCl, 0.4 mM MgCl2, and 0.42 mM CaCl2 were titrated to the given pH with NaOH. For reduced Cl− experiments, 133 mM NaCl was replaced with 133 NaGluconate, and MgCl2 and CaCl2 were raised to 0.74 and 1.46 mM, respectively, to account for gluconate-divalent binding. The amount of dilute HCl or NaOH added to medium to reduce pH to target level was determined empirically.

Respiratory capacity is a measure of the maximum rate of O2 consumption and mitochondrial electron transport in a cell. Glycolytic capacity is the maximum rate of glucose conversion to pyruvate and/or lactate by a cell. Glucose breakdown to two lactates produces two protons, allowing for the capability of indirect measurement of glycolytic rate using the extracellular acidification.

### Mathematical modeling of OVCAR3 growth dynamics

Growth dynamics were modelled with an ordinary differential equation (ODE) logistic model described by the following expression 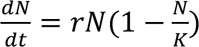, where N(t) is the population size at time t, r is the cell growth rate (in day^−1^), and K is the carrying capacity. Model parameters were estimated in Julia to estimate both r and k for the naïve and persistent populations individually under untreated and glutamine added media using the DifferentialEquations.jl [39] and DiffEqParamEstim.jl (https://www.github.com/SciML/DiffEqParamEstim.jl) packages. Cell counts were normalized by the first count read for each set of conditions to use for parameter estimation with an initial population size of 1. The ODEproblem function was used to define the problem and a loss function was created based on that ODEproblem to reduce the loss between experimental and simulated data using the Tsit5() solver and 10000 iterations. The optimized solutions returned the estimated growth rate (r) and carrying capacity (K) for the data for each population under each condition that are reported in **Supplemental Table T22.**

### Co-culture model and simulations of OVCAR3 growth dynamics

Based on the recurrence development in ovarian cancer, tumors are thought to be composed of heterogenous populations of cells. In our model, we assumed naïve and persistent cells compete in the tumor microenvironment following the equations below adapted from Strobl et al. [40]:

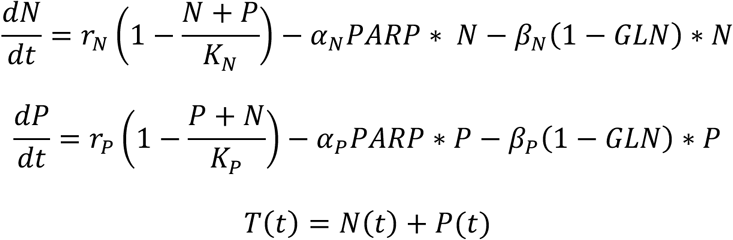

where N(t) and P(t) are the naïve and persistent population sizes at time t, *r*_*N*_ and *r*_*p*_ are the cell growth rate (in hours^−1^), and *K*_*N*_ and *K*_*P*_ are the carrying capacities for the naïve and persistent population previously estimated under normal conditions (0μM Olaparib/Gln+). The first term of both the naïve and persistent equations represents the logistic growth rate of the populations. In this case, the competition of the populations is modelled in a Lotka-Volterra fashion where the inter- and intraspecies competition coefficients both take the value of 1.

The second and third term of the equations represent the effect of treatment and glutamine deprivation in a linear manner assumed that this affected cell growth directly by a fixed amount. These terms were modelled as an on and off term that would change based on the magnitude of a factor of *α*_*N*_ and *α*_*P*_ for treatment effect and *β*_*N*_ and *β*_*P*_ for glutamine deprivation. Both the *PARP* and *GLN* took values of either 0 or 1. For instance, under untreated conditions the *PARP* term would take a value of 0 and not contribute to the growth of the population. Under treated conditions, the *PARP* term would take a value of 1 decrease population growth by a factor of *α*. Contrarily, when *GLN* is 0 the population would experience a decrease in growth by a factor of *β* since this term models the effect of glutamine deprivation.

*α*_*N*_ and *α*_*P*_ for 20μM Olaparib/Gln+ were estimated individually for naïve and persistent by fitting the mono-culture data for those conditions to the ODE equations above containing the *r*_*N*_, *r*_*p*_, *K*_*N*_ and *K*_*P*_ previously estimated under 0μM Olaparib/Gln+. These *α*_*N*_ and *α*_*P*_ were recorded as the effect of treatment under glutamine added conditions. A similar process was followed when estimating for *β*_*N*_ and *β*_*P*_ for 0μM Olaparib/Gln−. These were also parameterized for naïve and persistent individually by fitting the mono-culture data for those conditions to the ODE equations above containing the *r*_*N*_, *r*_*p*_, *K*_*N*_ and *K*_*P*_ previously estimated under 0μM Olaparib/Gln+. These *β*_*N*_ and *β*_*P*_ were recorded as the effect of glutamine deprivation under untreated conditions. To estimate all *α*_*N*_, *α*_*P*_, *β*_*N*_ and *β*_*P*_ under 20μM Olaparib/Gln− conditions, the mono-culture data for those conditions was used to fit the ODE equations above containing the *r*_*N*_, *r*_*p*_, *K*_*N*_ and *K*_*P*_ previously estimated under 0μM Olaparib/Gln+. These last *α*_*N*_, *α*_*P*_, *β*_*N*_ and *β*_*P*_ were recorded as the effect of treatment under glutamine-deprived conditions. All these parameters were estimated using the DifferentialEquations.jl [39] and DiffEqParamEstim.jl (https://www.github.com/SciML/DiffEqParamEstim.jl) packages in Julia. The ODEproblem function was used to define the problem and a loss function was created based on that ODEproblem to reduce the loss between experimental and simulated data using the Tsit5() solver and 10000 iterations. The effect of treatment and glutamine deprivation parameters are reported in **Supplemental Table T23**.

### Adaptive therapy schedules simulations

Growth rates, carrying capacities, glutamine deprivation terms and treatment terms previously determined for naïve and persistent populations were used within the co-culture ODE system to create adaptive therapy schedules simulations. To implement these adaptive therapy schedules, the differential equation model was modified at specified time intervals using Julia’s built-in Callback() function. This function allows for conditional triggers based on time or population state. For our simulations, treatment adjustments were made every 48 hours, with an initial drug dosage of 20 μM for all schedules. Two adaptive therapy treatment schedules were tested. The first adaptive therapy schedule (AT modulating) dynamically adjusted the drug dosage based on the tumor population size. If the tumor grew by 10% or more, the Olaparib dosage would increase by 5μM for the next 48-hour period. Conversely, if the tumor population decreased by at least 10%, the drug dosage would be reduced by 5μM. If the tumor population showed minimal change, the drug dosage would remain unchanged for the next interval. This approach aimed to simulate a physician adjusting treatment based on tumor progression. The second adaptive therapy schedule (AT skipping) employed a dose-skipping strategy. In this schedule, if the tumor population increased by 10% or more, the dosage would be set to 20μM. If the population decreased by at least 10%, the dosage would be reduced to 0μM. If there was not a net change of at least 10% in 48 hours the drug dosage in the simulation remained the same. Thus, the drug dosage alternated between two levels: 0μM or 20μM, based on tumor size.

Both adaptive therapy schedules were tested across 768 hours and treatment values were allowed range from 20μM to 0μM, sticking to either 20μM or 0μM for the AT skipping. To evaluate the effects of glutamine, for each adaptive therapy schedule both glutamine added, and glutamine deprived conditions were evaluated.

### Statistical analysis

The statistical tests were conducted in GraphPad Prism (Version 10.0.2). To analyze statistical differences between groups in CCK8 assay analysis, and between persistent and naïve populations’ pathway scores we used the non-parametric Mann-Whitney U test.

## ACKNOWLEDGEMENTS & FUNDING

Experimental research reported in this publication is supported by Physical Sciences Oncology Network at the National Cancer Institute through, Grant/Award Numbers: U01CA261841-01, and National Institutes of Health, Grant/Award Number: R01CA249016-01 for Dr. Damaghi’s team. The analysis research is supported by the UF Health Cancer Center, supported in part by state appropriations provided in Fla. Stat. § 381.915 and the National Cancer Institute of the National Institutes of Health under Award Number P30-CA247796. The content is solely the responsibility of the authors and does not necessarily represent the official views of the National Institutes of Health or the State of Florida. This work was in part supported by funding from the National Institutes of Health (NCATS 1KL2TROO1429). This work has been supported in part by the Molecular Genomics Facility Shared Resource at the H. Lee Moffitt Cancer Center and Research Institute, an NCI-designated Comprehensive Cancer Center (P30-CA076292).

## AUTHOR CONTRIBUTIONS

**A. Del Pino Herrera:** data curation, formal analysis, investigation, software, visualization, writing-original draft, writing-review & editing.

**J-H. Song, K. Torres-Acirga, S. Halder, J. Obeid**: performed experiments and analyzed the experimental data.

**K. El Bakkouri**: formal analysis, software, and visualization.

**W.M. Burke**: writing–review & editing.

**M. Damaghi:** MD conceived and designed the study, performed single cell experiments and analyzed the data as well as writing–original draft, writing–review and editing.

**M.C. Ferrall-Fairbanks**: data curation, formal analysis, investigation, methodology, project administration, resources, software, supervision, visualization, writing–original draft, writing–review & editing.

## DATA AND CODE AVAILABILITY

The code and documentation describing our single-cell analysis pipeline are open-source and publicly available through the OvCa PARP Persistence GitHub repository (https://www.github.com/mcfefa/OvCa-PARP-resistance). Three virtual machines producing the full computational environment is available on Code Ocean:

- R (https://www.codeocean.com/capsule/7415977/tree),
- Python (https://codeocean.com/capsule/2991364/tree),
- Julia (https://codeocean.com/capsule/6338854/tree)

